# Identifying transcription factor-DNA interactions using machine learning

**DOI:** 10.1101/2022.03.10.483780

**Authors:** Sohyun Bang, Mary Galli, Peter A. Crisp, Andrea Gallavotti, Robert J. Schmitz

## Abstract

Machine learning approaches have been applied to identify transcription factor (TF)-DNA interaction important for gene regulation and expression. However, due to the enormous search space of the genome, it is challenging to build models capable of surveying entire reference genomes, especially in species where models were not trained. In this study, we surveyed a variety of methods for classification of epigenomics data in an attempt to improve the detection for 12 members of the Auxin Response Factor (ARF) binding DNAs from maize and soybean as assessed by DNA Affinity Purification and sequencing (DAP-seq). We used the classification for prediction by minimizing the genome search space by only surveying unmethylated regions (UMRs). For identification of DAP-seq binding events within the UMRs, we achieved 93.54% accuracy, 6.2% false positive, and a 43.29% false negative rate across 12 members of ARFs of maize on average by encoding DNA with count vectorization for k-mer with a logistic regression classifier with up-sampling and feature selection. Importantly, feature selection helps to uncover known and potentially novel ARF binding motifs. This demonstrates an independent method for identification of transcription factor binding sites. Finally, we tested the model built with maize DAP-seq data and applied it directly to the soybean genome and found unacceptably high false positive rates, which accounted for more than 40% across the ARF TFs tested. The findings in this study suggest the potential use of various methods to predict TF-DNA interactions within and between species with varying degrees of success.

## BACKGROUND

Uncovering TF-DNA binding mechanisms and associated DNAs bound by TFs is important because of the impact gene expression has on phenotypic variation. One aspect that significantly modulates this process is the binding of transcription factors (TFs) (Cheng, et al., 2012). TFs bind to specific DNA sequences in the genome including promoters, enhancers and silencers to initiate, enhance or repress gene expression (Latchman, 1997). Alterations to TF-DNA binding sequences causes phenotypic variation by altering the levels of gene expression (Epstein, 2009; Pennacchio, et al., 2013). For example, in maize, the emergence of the enhancer for the *TEOSINTE BRANCHED (Tb1)* gene causes greater apical dominance due to higher *Tb1* expression compared to its ancient progenitor teosinte (Studer, et al., 2011). This particular domesticated DNA region for *Tb1* is located 65 kilobase pair (kbp) upstream and it functions as an enhancer (Bulger and Groudine, 2011; Studer, et al., 2011). The distal TF-DNA binding by enhancers make them challenging to detect compared to promoters that are located 50-100 base pairs (bp) upstream of the transcriptional start site (Siggers and Gorda^n, 2014). Moreover, the various patterns of TF binding make them difficult to predict compared to promoters that are bound by general TFs including TFIIB, TFIID and ^ RNAPII (Haberle and Stark, 2018).

Many experimental and computational techniques have been developed in an attempt to identify DNA regions where TFs bind. Chromatin Immunoprecipitation (ChIP) has been widely used to detect enhancers and silencers based on TF binding as well as chromatin modifications associated with DNA-bound TF (Huang, et al., 2019; Lu, et al., 2019; Oka, et al., 2017; Park, 2009; Ricci, et al., 2019). ChIP identifies DNA-interacting TFs by treating the cells with formaldehyde to crosslink TFs with DNA *in vivo*. Next, cells are lysed and chromatin is isolated and further fragmented. Crosslinked TF-DNA interactions are then captured by specific antibodies against the TF of interest. The frequency and strength of TF-DNA interactions are measured in a quantitative manner genome wide upon high-throughput sequencing, which is referred to as ChIP-seq. Although ChIP-seq is the gold standard for identifying TF-DNA interactions, it is difficult to perform and is especially challenging in species where antibodies are not easily obtained for performing immunoprecipitations of TFs of interest (Park, 2009). This limitation of ChIP-seq has led to the innovation of another technique to study TF-DNA interactions that is referred to as DNA Affinity Purification and sequencing (DAP-seq) (Bartlett, et al., 2017; O’Malley, et al., 2016). ChIP-seq captures DNA associated TFs *in vivo*, whereas DAP-seq identiies TF-DNA interactions *in vitro* (Bartlett, et al., 2017). For DAP-seq, fragmented genomic DNA with adaptors and affinity-tagged TFs are prepared separately (Bartlett, et al., 2017). TFs are then combined with the adapter-ligated fragmented DNA to allow for sequence-specific binding to genomic DNA *in vitro*, which is measured using high-throughput sequencing. As DAP-seq does not require a TF-specific antibody to capture TF-DNA complexes, it allows for screening of high numbers of TF-DNA interactions in comparison to ChIP-seq. Regardless, both methods have proven useful for the investigation of the genome-wide location of enhancers and silencers.

Computational approaches to predict TF-DNA interactions are actively being developed, despite the existence of experimental methods (Li, et al., 2018). The major driving force is that experimental methods are cumbersome and are not as scalable as computational methods. The most widely used computational approach is a supervised motif search using a position specific score matrix (PSSM) also known as position weight matrix (PWM) (Stormo, et al., 1982). Most motifs are ∼4-12 bp and PSSM builds the probability for the occurrence of each nucleotide at specific positions based on known TF binding motifs of interest (Mrázek, 2009). A sliding window-based approach is used where the window size is the size of the motif, and each sequence is scored against the PSSM to predict TF binding sites genome wide (Mrázek, 2009). As more motifs for diverse TFs are actively studied, this method can be expanded to predict regions bound by multiple TFs. However, as supervised motif searches do not provide accuracy about the search, it is challenging to identify functional TF-DNA bound regions. A major limitation is that this approach only finds sequences with a pattern match (Weirauch, et al., 2014). This leads to a high number of false positives, which misleads the characterization of true TF bound regions (Sielemann, et al., 2021). Moreover, motif based method searches can miss functional TF-DNA interactions for a number of reasons; 1) Some TF binding sequences do not have motifs (Inukai, et al., 2017), 2) Some TF binding sequences have multiple binding motifs (Nakagawa, et al., 2013), and 3) Recognition by TFs is also dependent upon sequences surrounding the motifs, so it is not enough to only identify the motifs alone (Inukai, et al., 2017).

Machine learning algorithms offer one possible solution for identifying the genome-wide landscape of TF-DNA interactions. Machine learning algorithms can learn and predict complicated patterns from data and provide accuracy about the prediction, a challenge that is suited for detection of TF-DNA interactions. In a study by Zamanighomi et al., motifs from DNA-binding domains were predicted with 60% accuracy using epigenomic data, although these models had relatively low specificity (Zamanighomi, et al., 2017). In contrast, a study by Mejía -Guerra et al. predicted TF-DNA interactions with a logistic linear algorithm and achieved >90% accuracy (Mejía -Guerra and Buckler, 2019). In this study, they used the flanking sequences around motifs to classify them into two classes, 1) TF binding sequences and 2) Non-TF binding sequences (Mejía -Guerra and Buckler, 2019). However, there are some limitations to these models; the models did not use the entire genome, but instead relied on simulated data. Lastly, Cochran et al. used deep learning to predict TF-DNA bound by several TFs, however, they reported high false positives (Cochran, et al., 2021). Collectively, these studies show the potential use of machine learning algorithms to identify functional TF bound regions, yet also highlight the challenging nature of this pursuit.

The development of experimental and computational approaches will enable the discovery of plant TF bound regions and their associated TFs important for gene regulation and phenotypic variation (Weber, et al., 2016). In this study, we use the well characterized Auxin Response Factor (ARF) family of TFs to build predictive models for detection of TF-DNA interactions. ARF TFs control target gene expression by responding to the plant hormone auxin and genome-wide maps of TF-DNA interactions has been generated in maize and Arabidopsis (Galli, et al., 2018; O’Malley, et al., 2016; Oh, et al., 2014; Ulmasov, et al., 1999; Wei, et al., 2021). As auxin plays a crucial role in plant growth and development, this TF family is an important test case for understanding our ability to predict TF-DNA interactions in plants (Li, et al., 2016). Here, we built a variety of models using ARF DAP-seq data from maize and applied some of them to the soybean genome to test their functionality. DAP-seq data for 12 members of the maize ARF TF family were tested with soybean genomic DNA to evaluate the validity of these models. Overall, our results show that count vectorization for k-mer and logistic regression models are effective at predicting TF-DNA interactions in maize, yet suffer from a high false positive rate when applied to the soybean genome. Collectively, this study shows the potential use of machine learning algorithms to identify TF-DNA interactions.

## RESULTS

### Preprocessing of data

To determine the ability to build machine learning (ML) models to predict TF-DNA interactions, we used previously published DAP-seq data from the maize ARF family (Galli, et al., 2018). We also performed DAP-seq using the same maize ARFs on soybean genomic DNA libraries. 12 ARF datasets were subsequently used for the analysis, as they had greater than a 1.5% FRiP (fraction of reads in peaks) score in both maize and soybean. Out of the 12 selected ARFs, six belong to clade A (ARF4, ARF16, ARF18, ARF27, ARF29, ARF34) and six to clade B (ARF7, ARF10, ARF14, ARF25, ARF36, ARF39). Clade A ARFs are known for their roles in transcriptional activation, whereas clade B ARFs are likely acting as repressors of transcription, in antagonism to clade A ARFs (Kato, et al., 2015; Kato, et al., 2020). Previous studies in Arabidopsis and maize reveal that clade A ARFs bind to the TGTCGG motif, whereas clade B ARFs prefer TGTC motifs with a cytosine tail such as TGTCCCCC (Boer, et al., 2014; Galli, et al., 2018). Because this TF family is so well characterized, it provides a unique opportunity to test the ability to build ML models to predict TF-DNA interactions across plant genomes.

For the 12 maize ARF datasets we selected to build ML models, there was an average of 37,840 binding events from the DAP-seq data, which accounts for 0.35% of the maize genome. This creates a unique challenge for machine learning models that rely on classification techniques, as the majority of the genome search space is devoid of ARF binding events. This leads to unbalanced data between bound and unbound regions and artificially inflates the prediction of classes with the higher numbers in the training datasets used. To generate a more balanced dataset, we limited the genome search space to Unmethylated Regions (UMRs), which are highly enriched for TF-DNA interactions compared to methylated regions in the maize genome (Crisp, et al., 2020). A previous analysis of UMRs in maize identified ∼100,000 regions, which accounted for ∼5.8% (123,146,800 bp) of the maize genome (Crisp, et al., 2020). A total of 11,149 ARF DAP-seq binding events overlapped a UMR, which accounted for 1.82% of the UMRs in maize (Supplementary Table 1). The percentage of ARF-bound regions increased about fivefold upon using UMRs compared to whole genome and helped to reduce the massive unbalanced data issue, however, 98% of the regions used in classifications are still unbound according to the DAP-seq data.

**Table 1.**
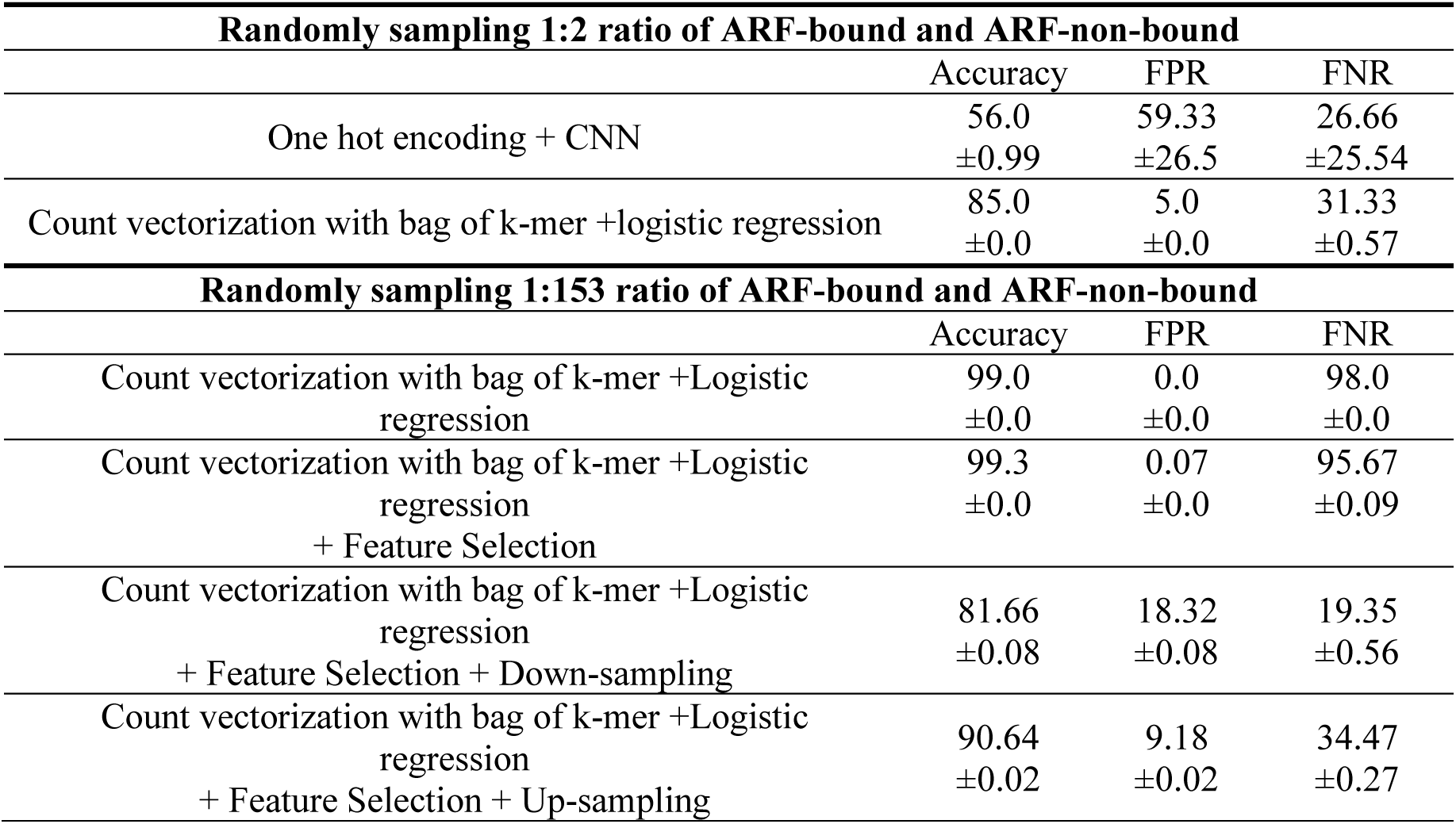
The performance of models using maize data from random sampling for 12 members of ARFs. Balanced data were used to evaluate two encoding methods (one-hot encoding and count vectorization with k-mer). To make the data, peaks from 12 member of ARFs were randomly picked to evaluate additional methods of classification that resulted in variable performance.

Unbalanced data leads to high accuracy of ML models, yet they are accompanied by a high false positive rate (FPR), as in this case ARF-bound regions would be falsely classified as ARF non-bound regions. This poses a challenge for evaluation of preprocessing of data, such as changes in bin sizes and labeling methods used for classification. Therefore, we used the same amount of input data for each class (ARF bound versus ARF non-bound) to find the optimal setting for data preprocessing. Because UMRs are longer than the DAP-seq peaks, we partitioned each subregion of a UMR into one of two classes (ARF bound and ARF non-bound) based on DAP-seq binding events (Figure 1). As with any protein-DNA enrichment-based sequencing assay, there is a distribution of sequenced fragments that decays over distance from the specific interaction site (summit), which is often attributed to sonication/fragmentation used in the assay. This produces regions that are challenging to classify as ARF bound or ARF non bound, which led us to classify the sequences flanking the summit as ‘ambiguous regions’. These regions were subsequently tested as ARF-bound or ARF-non-bound regions. We evaluated 75, 100, 125, 150, 175 and 200bp windows for classifications to identify the optimal window size for the analysis.

**Figure 1.**
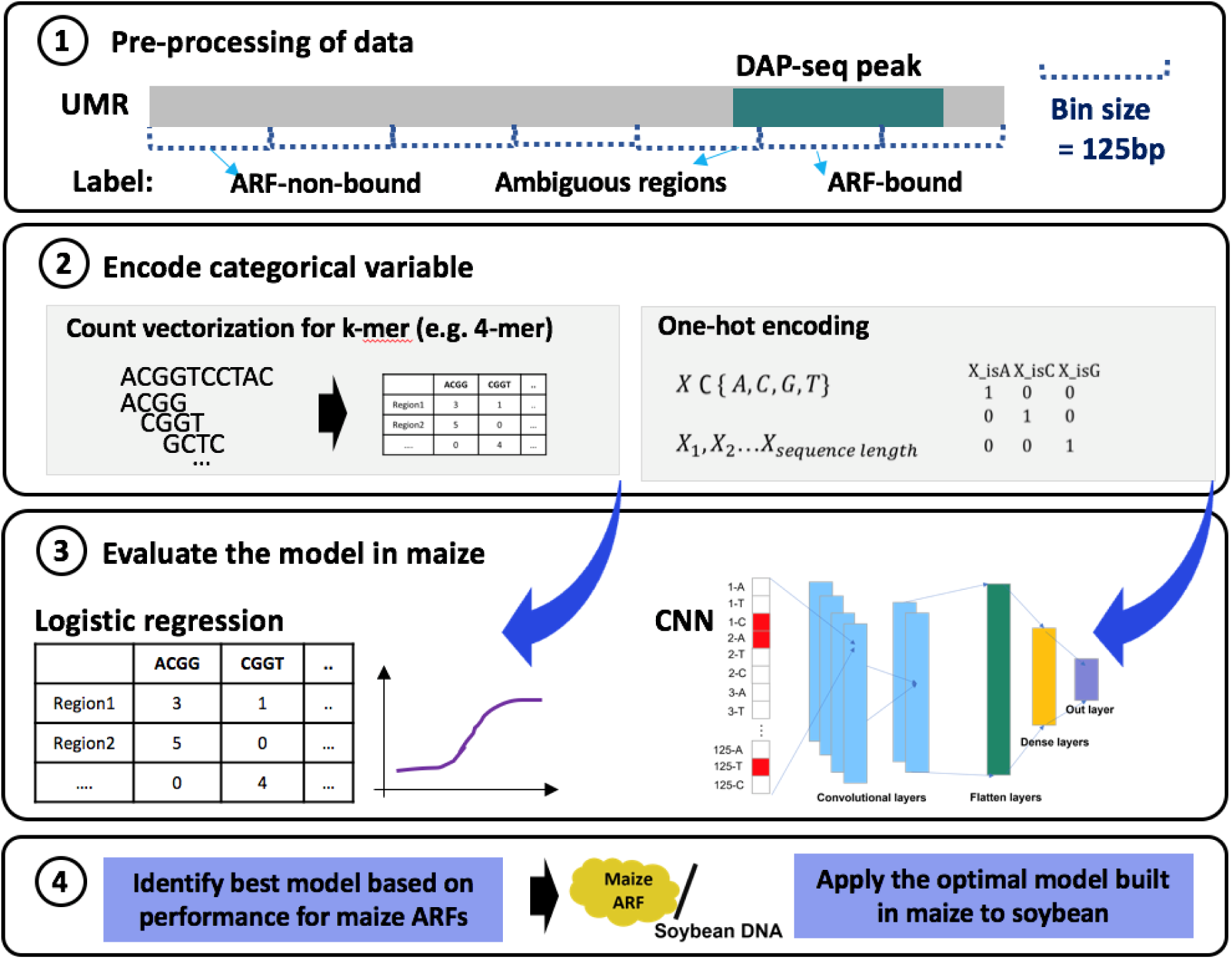
Experimental design and data processing. A diagram representing a whole experimental design for this research. After pre-processing of data, this research consists of two major steps for analysis: (1) Build the best model for 12 ARF members in maize; (2) Apply the model built by maize to soybean UMR.

We built models using all combinations of bin sizes and labelling methods using a grid search to the combination that produced optimal results. When ambiguous regions were considered as ARF-bound regions, the FPR was 24.49%, which is higher than the 6.73% observed when they were considered as ARF-non-bound regions (Supplementary Table 2). On the other hand, considering ambiguous regions as ARF-bound regions had higher false negative rate (FNR) compared to ARF-non-bound regions with ∼14% difference. As reducing the FPR is crucial to improve the classification performance from imbalanced data, we labelled ambiguous regions as ARF-non bound for subsequent analysis. Among the five bin sizes evaluated, the 125bp bin had the lowest FNR (18.87%). Thus, we found that partitioning (bin size=125bp) and labeling the ambiguous regions as ARF-non-bound regions were the most optimal for use in the next steps. This classification scheme resulted in a range from ∼1:50-1:700 (average 1:134) for the ratio of ARF-bound events to ARF-non-bound events for the 12 ARFs evaluated in this study (Supplementary Table 3).

### Building the best prediction models for identification of TF-DNA interactions in maize

Two distinct encoding methods such as one-hot encoding and count vectorization for k-mer have been used for DNA sequences for ML approaches (Yang, et al., 2020). We compared the two encoding methods with processed data with ARF bound and ARF non-bound in a 1:2 ratio to evaluate them without issues from imbalanced data. One-hot encoding implements the transformation of four nucleotides into binary information, that allowed us to apply a convolutional neural network (CNN) model as previously described (Quang and Xie, 2016). For the balanced data, one-hot encoding produced an accuracy of 56%, which was substantially lower than the 88.1% observed using count vectorization with k-mer and logistic regression. Thus, we applied count vectorization with k-mer and logistic regression for all subsequent analyses. To find the optimal length of the k-mer used to build models, we tested a range from 5-mer to 9-mer and ultimately selected to use a 7-mer, as it produced the lowest FNR of 31% (Supplementary Table 4).

The average number of events in each class for the 12 ARFs used in this study was 917,948.08 ARF- non-bound and 6,833 ARF-bound regions, which results in a 134:1 ratio. To reduce any effects due to individual ARFs, we randomly selected the average number of DAP-seq binding events (37,840) to produce the balanced data by random sampling, which resulted in a similar ratio (153:1) to the average number for ARF-non-bound and ARF-bound regions. The more imbalanced data in random sampling compared to the 1:2 ratio used above resulted in an increased FNR from ∼31% to almost 98%, which implies that almost all regions were classified into the ARF-non-bound class as this class is so dominant (Table 1).

One of the main factors that affects performance in ML is feature selection. Features are inherent properties of the data that ML models use to make predictions, as features distinguish different classes (Saeys, et al., 2007). We performed feature selection for two main reasons; 1) to improve the prediction performance by reducing the FNR and 2) to help identify features that are more likely to affect predictions. We used the frequency of 7-mer sequences within 125bp bin as features, as they potentially represent transcription factor binding sites. We also considered the pattern of features from the forward strand the same as the reverse strand, as ARFs recognize the patterns from both strands. This resulted in 8,192 (4 nucleotides^7^ divided by 2) features that were further reduced to 7,560 using entropy to eliminate features that have low complexity sequences (e.g. “AAAAAAA”). For feature selection, we filtered 2,222 features that had low variance in frequency across each 125bp window, as they do not provide enough information to help distinguish the window. After feature selection there were 5,338 features that we used to evaluate the effect of feature selection, which reduced the FNR to 95.67% from 98% (Table 1). This demonstrates that the model we built achieved a higher performance with a smaller number of features.

In addition to feature selection, we performed down-sampling and up-sampling to balance the training set and lower the false negative rate. Up-sampling increases the sample size for ARF-bound regions in the training data by randomly selecting greater numbers of ARF-bound regions whereas down-sampling decreases the sample size for non-ARF bound regions in the training data by randomly choosing less numbers of non-ARF bound regions. Down-sampling and up-sampling reduced the FNR to 19.35% and 34.47%, respectively (Table 1). The FPR was higher in down-sampling (18.32%) compared to up- sampling (9.18%). As the ARF-non-bound regions that are falsely classified into ARF-bound regions are reduced to 9.18% using up-sampling, we chose to implement the up-sampling method as it was the optimized method.

When the optimized method is applied to the 12 ARFs, an accuracy of 93.64%, a FPR of 6.2% and a FNR of 43% was observed on average across all TFs tested (Figure 2A). The high FNRs are due to the high number of ARF-bound regions that are falsely classified as ARF-non-bound regions. Clade A had a FNR of 28%, which is lower than the FNR of 57.7% observed for clade B. Considering that clade A has a higher number of ARF-bound regions on average from the DAP-seq data, this results in less imbalanced class numbers compared to clade B, which likely explains the differences in the FNRs. A significant correlation of the ratio for the two classes and the FNR was observed, which demonstrates that the imbalanced data affects the FNR (Figure 2B). In contrast, the FPR showed a negative correlation with the greater number of imbalanced classes (Figure 2B). The FPR is calculated by dividing the FP by sum of the FP and the TN. When the data are more imbalanced with a greater number of ARF-non- bound regions, a greater number of ARF-non-bound regions (Negative) are classified as TN or FN. This explains the negative correlation observed between the FNR and the FPR (Figure 2B); ARF34 had the lowest FNR (18.1%) and the highest FPR (12%), whereas ARF39 had the highest FNR (74.52%) and the lowest FPR (1.14%) (Figure 2C).

**Figure 2.**
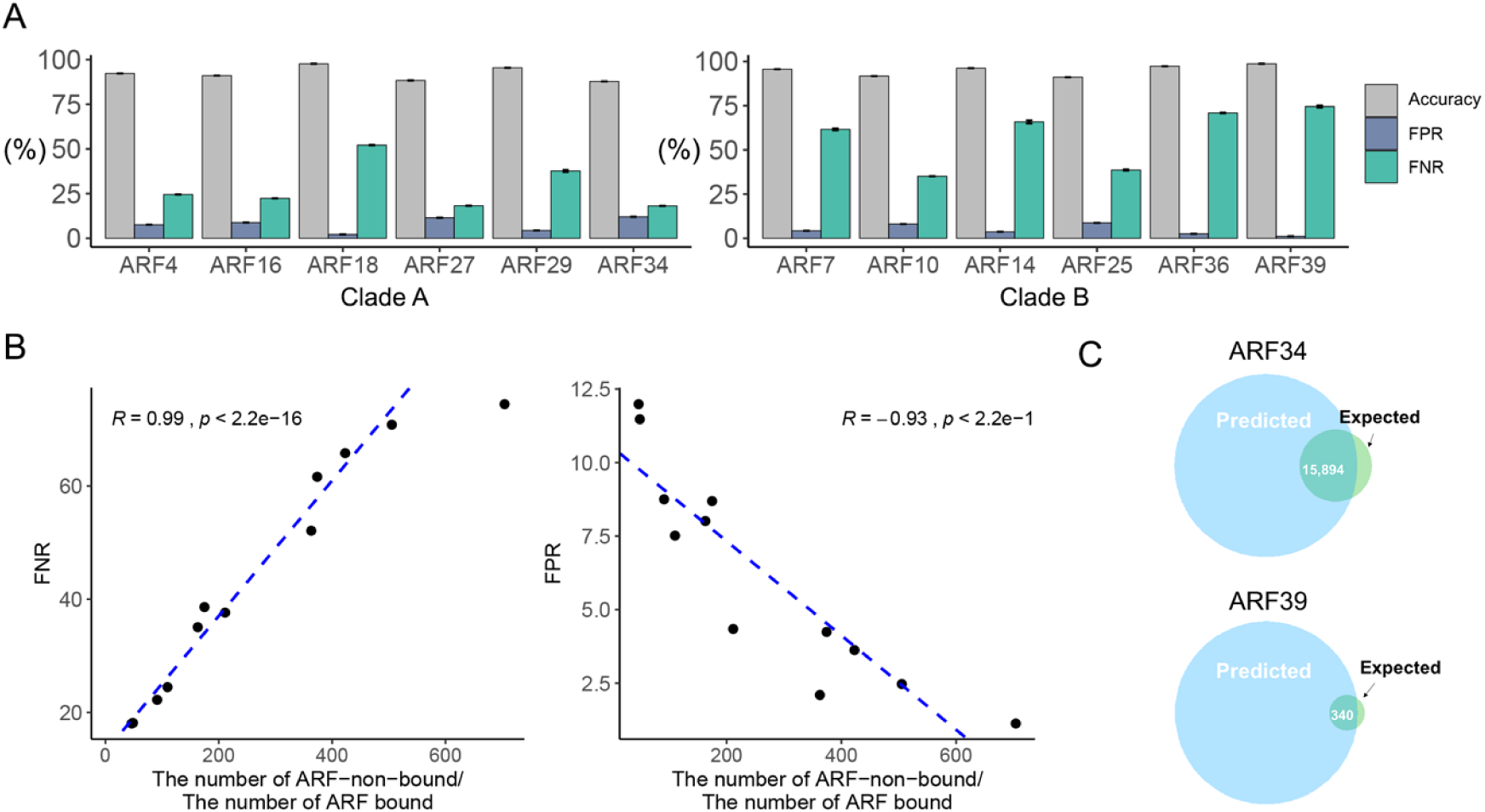
The performance of classification on prediction of binding for 12 ARFs from maize. (A) The accuracy (ACC), FPR and FNR for the classification of maize genome. (B) The correlation between FNR and the ratio of ARF non-bound/bound regions. Each dot represents an ARF that was tested. (C) Venn diagrams of the predicted number of TF-DNA interactions and the empirical results determined using DAP-

### Evaluation of features

The features used to build the logistic regression model distinguished ARF-bound and ARF-non bound regions in the maize genome with high accuracy. Even though the motif information is not used in the classification, the features used to build the predictions are expected to find the pattern of sequences where the ARFs are more likely to bind. To find which features negatively or positively affected the prediction, we investigated how much each feature affects the performance of the prediction. We defined the feature by log transformation of the coefficients of the logistic regression model for each of the ARFs. Selected features with high or low scores are indicative of genomic sequences where ARFs are more or less likely to bind. As the motif sequences for clade A and clade B from DAP-seq peaks are distinct in terms of the tails of cytosines (Figure 3A), we calculated the feature score individually for each clade. We identified the top 15 features with highest or lowest feature score out of the 5,338 total features (Figure 3B).

**Figure 3.**
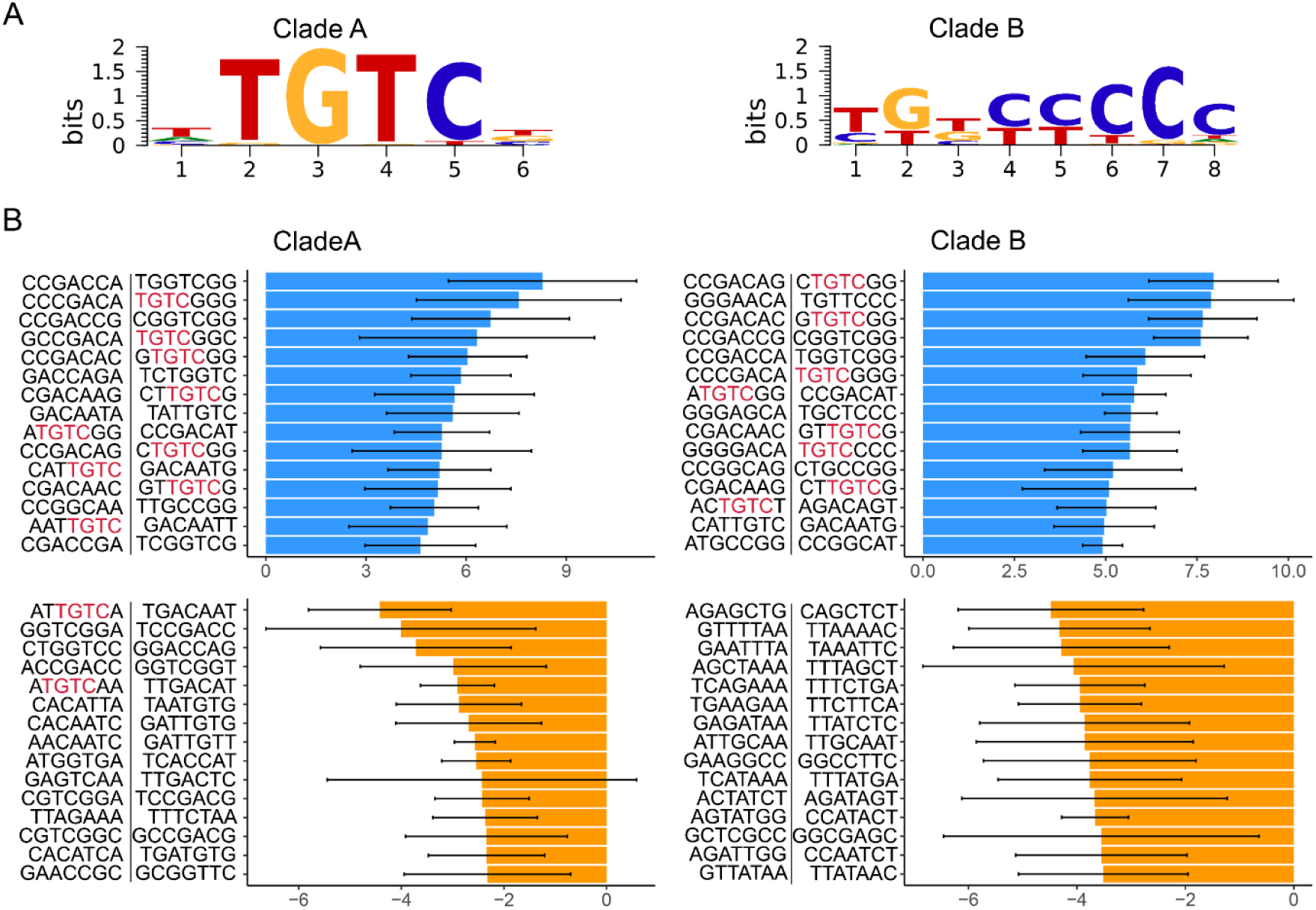
(A) Top motif identified for maize ARF clade A and clade B by identifying the combined peaks from clade A and clade B DAP-seq data. (B) The features with highest score or lowest score. The bar represents the average of feature score, and error bar represents standard deviation among ARFs in clade A or B. The upper box shows features with the top 15 highest feature score while the lower box shows features the prediction whereas “A” tails negatively affect the prediction. The features that have negative effects provides clues about which sequences ARFs do not prefer.

In clade A, “TGGTCGG”, “TGTCGGG”, “CGGTCGG”, “TGTCGGC” and “GTGTCGG” were the most important for model performance. Out of the top 15 features with the highest score, nine features contained the known ARF “TGTC” motif. Most of the features that have “TGTC” had a tail of “G” in clade A. The top five features identified in clade B were “CTGTCGG”, “TGTTCCC”, “GTGTCGG”, “CGGTCGG” and “TGGTCGG”. In clade B, out of the top 15 features with the highest score, eight features had “TGTC”; one with “C” tail and the other with “T” and six with “G” tails. The high feature scores for “TGTC” followed with “G” are not consistent with the motif sequences from DAP-seq peaks. Motif sequences from peaks have no following nucleotides in clade A and “C” tails in Clade B. The top 15 features in clade A and B with the lowest scores showed various patterns of 7-mers of nucleotides. Two of them in clade A contained the “TGTC” sequences followed with “A”. It is expected that the nucleotide that follows “TGTC” has an important role in the prediction; “G” tails can positively affect It was unexpected that the features with “TGTC” with “G” tails have positive effects on the classification, considering that maize ARF motifs from PWM had “TGTC” with “C” tails. To find if the ARF-bound regions have abundant features with “G” tails that affects the classification, we examined the frequency of features in ARF-bound regions. In clade A, “TGTCGGC”, “TTGTCGG”, “GCTGCTG”, “CTGTCGG” and “CTGCTGC” were the top 5 most abundant features across clade A. In clade B, features of “CTGCTGC”, “GCTGCTG”, “TGCTGCT”, “TGTCGGC” and “CTGTCGG” had dominant frequencies. They had “G” or “GG” sequences after “TGTC”. This implies that there are some binding sites that are overlooked by the PWM approach. This is one advantage of our approach to extract features from classifications for motif identification, as traditional motif detection methods do not provide sequence information for regions that have negative relationships with peaks.

### Evaluation of the efficacy of using the TF-DNA prediction models built using maize on the soybean genome

Many DNAs-bound TF are predicted to be conserved across evolution if they are important for conservation of specific traits or responses to the environment. This is especially true of the ARF gene family, as they have a conserved N-terminal DNA binding domain and in most cases a conserved C-terminal dimerization domain across plant species (Tiwari, et al., 2003). In Arabidopsis, ARFs bind to the “TGTCTC” motif and some ARFs that are important for transcriptional activation show a preference for binding the “TGTCGG” motif (Freire-Rios, et al., 2020). Additionally, the promoter regions for auxin-responsive genes in soybean are enriched for the “TGTCTC” motif (Guilfoyle, et al., 1998). Thus, we hypothesized that the model we built using the maize genome to predict TF-DNA interactions could be applied to the soybean genome. Considering maize is a monocot and soybean is a dicot, which have significant time since divergence (Chaw, et al., 2004), if the model can successfully predict TF:DNA interactions in soybean it would provide strong evidence that this model can be used to robustly predict TF:DNA interactions in other species that lack experimental data.

To test this hypothesis, we produced DAP-seq data for the same maize ARFs used in the first part of this study using soybean genomic DNA as input. To evaluate the quality of the DAP-seq data we produced using soybean genomic DNA, which is referred to as ZmARF to distinguish them with the ARF binding events in the maize genome. We investigated sequence alignment rates, peak shape and the fraction of reads in peaks (FRiP) at the 0.00001 FDR threshold, as we did for the maize DAP-seq data (Schmitz, et al., 2021). The DAP-seq libraries for the 12 ZmARFs showed more than 95% of alignment rates (Supplementary Table 5). Mapped data showed strong peak signals with low background noise (Supplementary Figure 1 and Figure 4A). Collectively, these DAP-seq experiments had a FRiP score of 10.59% on average with a range from 1.6-24% (Figure 4B). All ZmARFs except ARF18 had greater than a 2% FRiP score. Using the newly produced DAP-seq data, we identified ARF-bound regions and identified sequence motifs for all 12 ARFs tested to evaluate conservation of motif preferences (Figure 3B). In maize, the ARF binding motif predominantly clusters based on evolution of the ARF gene family (Galli, et al., 2018). The Pairwise Pearson correlation showed that binding profiles clustered according to their clade A or B phylogenetic classification (Supplementary Figure 2). However, the motifs identified using the maize ARFs screened against soybean genomic DNA showed that clade A ARF motifs (ZmARF4, ZmARF16, ZmARF18, ZmARF27, ZmARF29 and ZmARF34) clustered together, whereas clade B ARF motifs were distributed across the tree (Figure 4C). For example, ZmARF25 is a member of clade B, however it groups with clade A. The top enriched motif in the ZmARF7- and ZmARF10-bound soybean genomic DNA datasets were more diverse compared to other ZmARFs, although the reason for this is unknown at this time. Lastly, the binding motifs detected for clade B in soybean did not possess the long tail of “C” that is common in maize. It’s actually consistent with a previous result that investigated maize clade B ARF binding to Arabidopsis DNA (Galli, et al., 2018).

**Figure 4.**
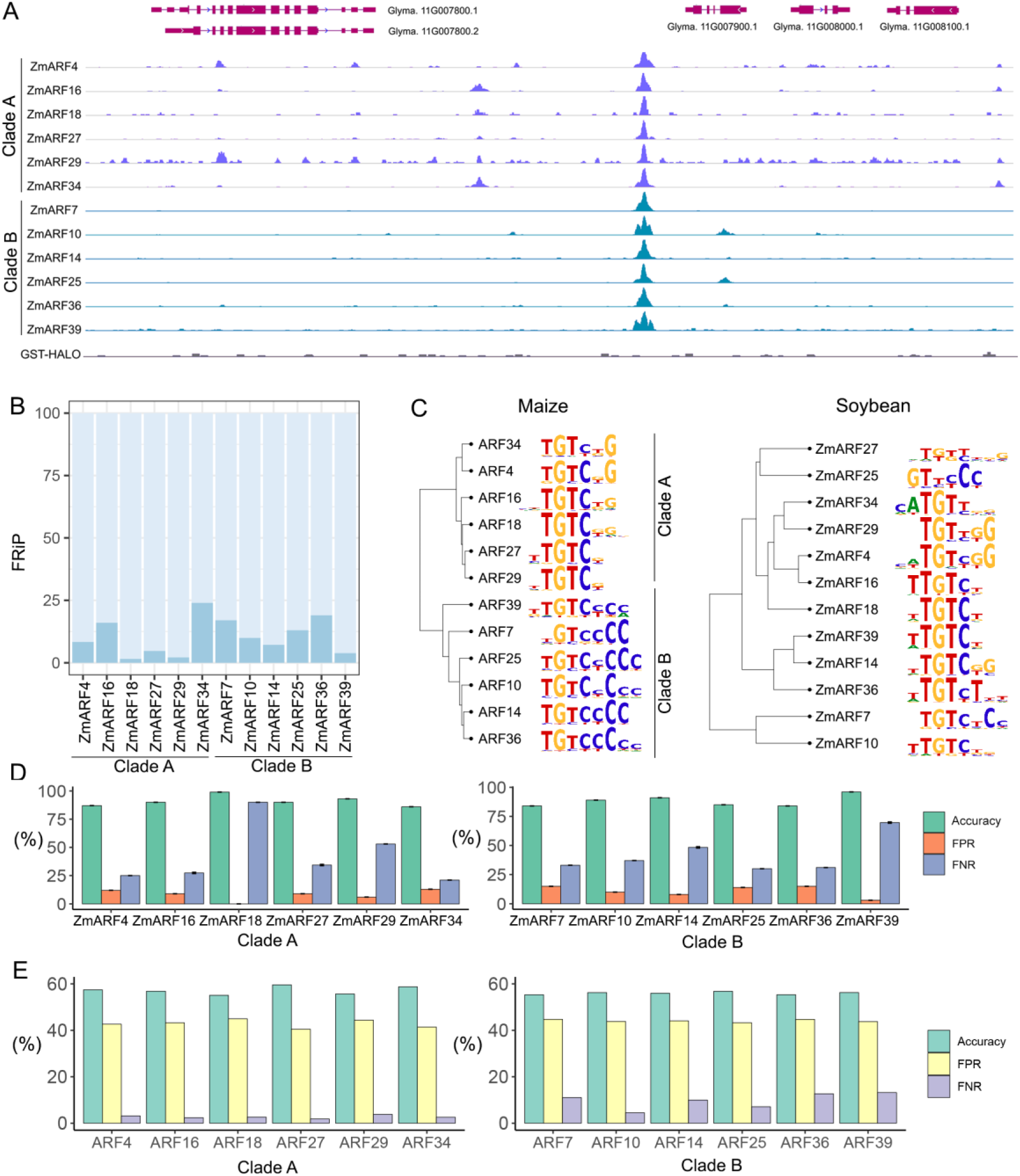
(A) ZmARF DAP-seq peaks in soybean. (B) FRiP (fraction of reads in peaks) of soybean DAP-seq data. (C) Dendrogram of the top maize and soybean ARFs based on profiles of the binding motif sequences. (D) The prediction performance of the model when it is built and evaluated against soybean data. (E) The prediction performance when we build the model with maize data and apply it to soybean data.

We used same data preprocessing methods we used for maize and applied them to the soybean DAP-seq data. In maize, the use of UMRs reduced the data imbalance issue as fivefold more DAP-seq peaks were located in UMRs compared to the entire genome. In soybean, the use of UMRs actually resulted in a small increase in the overlap with DAP-seq binding events (0.35%) compared to ratio observed across the genome (0.3%) (Supplementary Table 6). This is in large part due to the fact that soybean genome is less methylated than the maize genome. Nonetheless, we divided each UMR into 125bp bins and labelled them using the same method as we did for maize. Approximately 2,325,855 ZmARF-non-bound regions and 3,185 ZmARF-bound regions were identified for each ZmARF tested on average (Supplementary Table 7). Compared to the maize data which showed 1:134 ratio for bound vs unbound regions, soybean was significantly more imbalanced showing a 1: 1,575 ratio. We used the same model with maize by training and testing with soybean DAP-seq data (Figure 4D). Some ARFs such as ARF18 and ARF29 showed over a 50% FNR, which implies that the high number of ARF-bound regions are falsely predicted as ARF-non-bound regions. These results are somewhat consistent with what we observed in maize, if we consider that the soybean data has a significant imbalance due to a minority of ARF-bound regions, which is associated with a high FNR in both experiments (Supplementary Figure 3). Regardless, the soybean DAP-seq data was tested using the most optimal prediction model used for the maize ARF-binding data, which universally showed that the accuracies of predictions were below 59% with greater than a 55% FPR (Figure 3E).

## DISCUSSION

In this study, we compared the performance of classification in terms of the encoding methods and classification algorithms such as one-hot encoding with neural network algorithms and k-mer vectorization with logistic regression. Previous studies used one-hot encoding and neural network algorithms for genomic sequences and achieved high performance to identify TF binding motifs (Alipanahi, et al., 2015; Kelley, et al., 2016). Moreover, one-hot encoding with neural network algorithms showed high performance for classifying TF binding sequences (Mejía -Guerra and Buckler, 2019). In this study, one-hot encoding with neural network algorithms showed lower performance than k-mer vectorization with logistic regression. Although previous studies using one-hot encoding with neural network algorithms found the motif sequences among TF binding regions by predicting binding scores (Alipanahi, et al., 2015), this study used classification methods to classify the long length of the sequences into TF-bound sequences or TF non-bound sequences. In classification, k-mer scans ∼100bp sequences with a small unit of length to identify the specific sequence that TFs bind, whereas the one-hot encoding method recognizes the sequences as whole images. This implies that classifying ∼100bp sequences requires features that can specify distinguishing characteristics. This creates some limitations, especially for shorting binding motifs within the 100bp input windows, that could be mitigated using additional inputs such as using accessible chromatin regions that are enriched for TF binding (Kelley, et al., 2016).

We built a model using 12 members of ARF TF gene family in maize and evaluated the prediction performances between clade A and B. Although we showed that the FNR and the FPR are correlated with the ratio of number of imbalanced classes, there are still other factors that can affect the performance. The differences of the performance between the clades were higher than the differences between ARFs within a clade. We expected that long tale of “C” in clade B should provide an advantage using k-mer vectorization, because the tail of Cs included in the 7-mer provides greater numbers of distinct characteristics of features. Furthermore, clade A binds to tandem repeat in auxin response elements, which can make finding binding events more difficult (Chandler, 2016). However, clade B showed lower performance compared to clade A in terms of the FNR. Therefore, the evaluation of features showed that the “C” tails after “TGTC” did not improve the performance, as the features with high impact had “G” tails not “C”. The unexpected result of the G tails could be due to the 7-mer length of the feature we used or result from the high FNR. Furthermore, the FNR and FPR are more affected by the imbalanced number of classes. Collectively, this implies that data imbalance influences the prediction performance more than the structure of motifs does.

Classification of data with imbalanced class distribution is well known to negatively impact performance (Estabrooks, et al., 2004). It is well established the majority of eukaryotic genomes are comprised on non-coding DNA sequence, a subset of which includes TF bound DNA sequences (Elkon and Agami, 2017). This feature of eukaryotic genomes leads to an unbalanced data issue, as the ratio of non- TF bound DNAs is much higher than that of TF bound DNA. Our research shows that the prediction comparing the data with balanced data and unbalanced data showed that imbalanced data can increase the FNR to 98% from 31.33%. The high FNR suggests that ARF-bound regions are falsely classified as ARF-non-bound regions. The algorithm in training steps recognized that the number of ARF-non-bound regions are more abundant than ARF-bound regions, thus it causes incentives of classifying the samples to ARF-non-bound regions. To reduce the drawback from the imbalanced number of classes, we added more weight to the minority class (ARF-bound regions) in the logistic regression, however, this did not show an improvement in classification performance (Li, et al., 2010). Subsampling for class imbalances, including down-sampling and up-sampling, were performed (Sun, et al., 2009). Even though this led to a reduction of the FNR using up-sampling, it increased the FPR by falsely classifying the ARF-non-bound regions as ARF-bound regions. The up-sampling methods makes algorithms more likely to classify samples to ARF-bound regions in the training set, but in the test set the true number of ARF-bound regions is much lower than ARF-bound regions. This implies that implementing classification algorithms has some drawbacks to identifying TF-DNA interactions given that most regions of genomes are not bound by TFs.

We validated the model established with maize against the soybean genome to determine if the model can be used to robustly predict TF-DNA interactions in other plant species. As ARFs share similar motif sequences between plant species (Tiwari, et al., 2003), we hypothesized that the model built using maize ARF DAP-seq data would predict ARF binding regions successfully. However, the application of the model to a different plant species showed high FPRs and low accuracies compared to the model tested with the same species. It is possible the divergence time between maize (monocot) and soybean (dicot) is too large preventing cross species application of these models (Chaw, et al., 2004). We found two main differences between maize and soybean DAP-seq data; 1) Motif shape and 2) The ratio of class number of ARF-bound and ARF-non-bound regions. Some ARFs such as ZmARF25, ZmARF27, ZmARF29 and ZmARF34, when tested with soybean genomic DNA did not show enrichment for binding the core motif of “TGTC”. Moreover, some members shared the same core motif of “TGTC”, but the sequences around “TGTC” were different. For example, ZmARF36 tested with maize genomic DNA had “C” tails after “TGTC” but when tested with soybean genomic DNA had “T” tails. Furthermore, the expanded proportion of UMRs present in soybean compared to maize led to greater imbalanced data when trying to predict ZmARF-bound regions in the soybean. It is assumed that the different distribution of ARF binding events between maize and soybean lead to an ‘Out of Distribution’ effect (Arjovsky, 2020). The model was designed to learn generalizable knowledge from the maize training data and it expected that the soybean test data would share the same distribution with the maize training data. This implies that the data distribution between the soybean and the maize genome are sufficiently different enough making it challenging to apply the same prediction model. Although this study shows the limitation of application to different species it is possible it could be improved in the future by focusing on more closely related species. Regardless, this study presents a unique approach and demonstrates the potential use of machine learning algorithms to identify TF-DNA interactions in plant genomes.

## MATERIALS AND METHODS

### preparation and DAP-seq

Genomic DNA libraries for soybean were prepared following the protocols in Bartlett et al(Bartlett, et al., 2017). Genomic DNA (gDNA) was extracted from leaf tissue using phenol:chloroform:IAA extraction. Five micrograms of gDNA was diluted in EB (10 mM Tris-HCl, pH 8.5), sonicated to ∼200 bp fragments in a Covaris S2 sonicator and purified with AmpureXP beads. Samples were end repaired using an End-It kit (Lucigen) and purified with AmpureXP beads. Purified samples were A-tailed using Klenow 3–5′exo- for 30 min at room temperature and then purified with AmpureXP beads. A Y-adapter was ligated as described in Bartlett et al. To attach the protein to the MagneGST beads(Promega), 20 μl of purified GST-ARF protein (5–20 μg) was diluted in 400 μl of 1X PBS containing 25 μl of washed beads. In addition to the GST-ARF samples, a negative control GST-HALO sample was performed using protein expressed in the TNT wheatgerm expression system (Promega). Beads were washed four times in 1X PBS + NP40 (0.005%) and resuspended in 100 μl of 1X PBS. 1μg of gDNA library was diluted to a final volume of 60 μl in 1X PBS and added to the protein bound beads. One microgram of genomic DNA library was diluted to a final volume of 60 μl in 1X PBS and added to the protein bound beads. Samples were then incubated for 1 hour at room temperature. Beads were washed in 1xPBS + NP40 and recovered by resuspending in 25 μl EB and boiling. Eluted samples were enriched and tagged with dual indexed multiplexing barcodes by performing 20 cycles of PCR in a 50 μl reaction51. We sequenced samples on a NExtSeq500 with 75 bp single end reads. A total of 10– 30 million reads were obtained for each sample. Maize DAP-seq data were downloaded from GEO accession GSE111857 produced by *Galli, M. et al*. (Galli, et al., 2018).

### Peak calling for DAP-seq data

The raw reads were trimmed by filtering out adaptor-only nucleotides with the following parameters ILLUMINACLIP:TruSeq3-SE:2:30:10 LEADING:3 TRAILING:3 SLIDINGWINDOW:4:15 MINLEN:50, using Trimmomatic (ver 0.36; (Bolger, et al., 2014)). Trimmed reads were aligned to the reference genome (Gmax505 v4.0) using bowtie2 v2.2.853(Langmead and Salzberg, 2012). Mapped reads with >MAPQ30 were removed to use the reads that are not mapped into multiple locations. We called the peaks using GEM v2.554 using the GST-HALO negative control sample with following parameters: --k_min 6 --k_max 20 --outNP –sl --q 5 (Guo, et al., 2012). The samples with more than 2% of FriP at the 0.00001 FDR threshold using ChIPQC v1.8.2 were chosen in all subsequent analysis(Carroll, et al., 2014). We used motifs discovered by GEM for the first round of motif prediction (Guo, et al., 2012). We represented sequence logos and dendrogram for motifs for the ARF family using motifStack based on a position count matrix (PCM) of the motifs (Ou, et al., 2018). The heatmap for binding events from Pearson correlation was calculated with 10bp bin size using deepTools v 3.5.1 (Ramírez, et al., 2014). For generation of metaplots, the signal densities for DAP-seq data were calculated with deepTools v 3.5.1 using the following parameters: ‘-a 3000 -b 3000 -bs 10’.

### Identification of UMRs

For UMRs in maize, previously identified maize UMRs (Crisp, et al., 2020) were used. UMRs in soybean were identified by followed steps. Two replicates of WGBS data for G. max Williams82 (pooled leaves) were downloaded from SRA, which are SRR12494495 and SRR12494496 (Wang, et al., 2021). Replicates were filtered and mapped individually, then the final mapping files (bams) were merged to increase coverage for UMR identification. Unmethylated regions (UMRs) were identified as per (Crisp, et al., 2020). Reads were trimmed using Trim galore! version 0.6.4_dev, powered by cutadapt v1.18 (Martin, 2011) and quality checked using fastqc v0.11.4. Next, 20 bp was trimmed from the 5’ ends of both R1 and R2 reads and aligned with bsmap v2.74 (Xi and Li, 2009) to the soybean v4 genome (Gmax_508, phytozome v13) with the following parameters -v 5 to allow 5 mismatches, -r 0 to report only unique mapping pairs, -p 1, -q 20 to allow quality trimming to q20. Conversion efficiency of 99.5% was determined by appending the chloroplast genome (NC_007942.1) to the v4 reference genome for mapping. Output SAM files were parsed with SAMtools (Li, et al., 2009) fixsam, sorted and indexed. Picard MarkDuplicates (v 2.9.0-1) was used to remove duplicates, BamTools filter (v 2.4.0) to remove improperly paired reads and bamUtils clipOverlap (v 1.0.13) to trim overlapping portion of paired-reads so as to only count cytosines once per sequenced molecule. The methylratio.py script from bsmap v2.74 was used to extract per-site methylation data summaries for each context (CH/CHG/CHH) and reads were summarised into non-overlapping 100bp windows tiling the genome. WGBS analysis pipelines are available on github (https://github.com/pedrocrisp/crisplab_epigenomics/tree/master/methylome). To identify unmethylated regions, each 100bp tile of the genome was classified into one of six domains or types, including “missing data” (including “no data” and “no sites”), “High CHH/RdDM”, “Heterochromatin”, “CG only”, “Unmethylated” or “intermediate”, in preferential order as per (Crisp, et al., 2020).

### Producing combined ARF data by random sampling

To test the bin lengths selected and the labelling method used, we produced random sampling data, which combined data from 12 members of the ARF gene family in maize. For random sampling data, we randomly selected the average number of peaks (37,840) from peaks from 12 members of the ARF gene family. Next, we performed data preprocessing by dividing the genome into various bin sizes and annotating them as ARF-bound or ARF-non-bound. This produced an unbalanced data set. Redundant regions were removed. Moreover, to produce a balanced random sampling data set, we randomly selected the same number of ARF-bound in ARF-non-bound regions and assigned half of the ARF-bound regions to ambiguous regions.

### Cross validation

We performed 5-fold cross validation for all predictions by dividing the data into five subsets. We used four data subsets (80% of data) for training and one data subset (20% of data) for testing with shuffling the subsets. We repeated it three times to show how the models are generalize dto different combinations of data sets.

### Features for Count vectorization of k-mer

To vectorize a sentence in natural language processing, bag of words can be applied(Zhang, et al., 2010). Bag of words counts the number of the occurrence for each token and uses the vectorization of counts information for training. Bag of words does not consider the order of words in the sentences. The genome sequence is read with a k size of sliding windows that is called k-mer. In this case, k is read the length of the word. For example, when there is a group of sequence of AATTG, tokens of 3-mer is AAT, ATT, TTG and TGC. To find the optimal length of the k-mer, we tested from 5-mer to 9mer and chose the length with the lowest FNR. We created sequences that contained the sequence of the feature as well as its complementary sequence, which were separated with an “N”. For example, for a 125bp window with 7-mer, which was labelled as an ARF-non-bound region, the possible feature sets used looked like the following sequences (“AATTGTTNAACAATT”:2, “AATTGGCNGCCAATT”:1,.., “CCCATACNGTATGGG”:1). In natural language processing, stop words such as “the”, “a”, “an” and “in” are filtered while tokenization. DNA sequences can have stop words as repeated sequences occur in multiple copies throughout the genome. The stop words for DNA were defined as the DNA sequences with low entropy. Thus, the feature with low entropy means the feature have low variances within the feature. In the previous studies, the repetitive regions are expected to carry little information for TF-DNA binding and low entropy was calculated by adding the probability of appearance of the i-th base in the token as the equation below(Mejía -Guerra and Buckler, 2019).

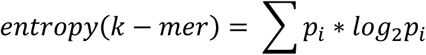

The tokens with lower than 1.3 entropy were considered as stop words according to the TF-DNA binding database and eliminated. We normalized the frequency of each feature so that all data were on the same scale for calculation of the variance. We used “scaler.fit_transform” to standardize the values for features with a standard score. For feature selection, features with low variance were eliminated using “VarianceThreshold” with a 0.001 threshold.

### One-hot encoding

One-hot encoding can be used to transfer DNA sequences to binary information. Then, learning algorithm such as Deep Neural Networks and Convolutional Neural Networks can be adapted to DNA by considering DNA as a fixed length 1-D sequence with four channels (A,T,G,C)(Alipanahi, et al., 2015). A, C, G, T will be encoded into (1 0 0), (0 1 0), (0 0 1), (0 0 0) respectively. For example, when the sequence is ATTGC, then it will be transformed to ((1 0 0), (0 0 0), (0 0 0), (0 0 1), (0 1 0)). As we use a length of 125 A,T,G,C sequences, the input data will have a 3-D structure with 3*125*the number of samples. Subsequently the 3-D data structure is flattended using ‘model.add(Flatten())’

### Down-sampling and Up-sampling

Down-sampling and Up-sampling is re-sampling techniques for training data to balance the training set and relieve the imbalanced data issue. Down-sampling randomly subsets samples from the class that has the dominant number of samples to match the least prevalent class(Estabrooks, et al., 2004). As we have more number of ARF-non bound regions than ARF bound regions, down-sampling randomly removed some ARF-non bound regions in training set to be matched with the same number of ARF-bound regions in training step. Up-sampling randomly samples the minority class to be the same size as the majority class in the training set(Estabrooks, et al., 2004). We increased numbers of ARF-bound regions in training set by randomly adding ARF-bound regions with the same number of ARF non-bound regions. In the test data set, the imbalanced data was used.

### Parameters for the models

For logistic regression we used the ‘LogisticRegression’ function in scikit-learn with L2 regularization penalty, 1e-4 tolerance, 1.0 C, Liblinear optimization and binary classification. To create the sequential model we used “tf.keras.models.Sequential” that can create linear sequences of processing layers with 10 epochs, 50 batch sizes, 0.1 validation split, 1 verbose. We used the function of Sequential() and add the layers with Conv1D() and MaxPooling1D() for 10 epochs, 50 batch sizes. To use 3D data structure of DNAs, we used the function of Flatten().

## DECLARATIONS

### Authors’ contributions

All authors were involved in this experiment, drafting the article or revising it critically for important intellectual content.

### Consent for publication

Not applicable

### Availability of data and material

Raw sequencing data are available at the NCBI (GSE193400).

## Acknowledgements

We would like to thank Andrea Brautigam for helpful suggestions to improve the quality and clarity of this study. We would also like to acknowledge support from the Georgia Advanced Computing Resource Center as well as the Georgia Genomics & Bioinformatic Core.

## REFERENCES

Alipanahi, B., et al. Predicting the sequence specificities of DNA-and RNA-binding proteins by deep learning. Nature biotechnology 2015;33(8):831–838.

Arjovsky, M.: New York University; 2020. Out of distribution generalization in machine learning.

Bartlett, A., et al. Mapping genome-wide transcription-factor binding sites using DAP-seq. 2017;12(8):1659.

Boer, D.R., et al. Structural basis for DNA binding specificity by the auxin-dependent ARF transcription factors. 2014;156(3):577–589.

Bolger, A.M., Lohse, M. and Usadel, B. Trimmomatic: a flexible trimmer for Illumina sequence data. Bioinformatics 2014;30(15):2114–2120.

Bulger, M. and Groudine, M. Functional and mechanistic diversity of distal transcription enhancers. Cell 2011;144(3):327–339.

Carroll, T.S., et al. Impact of artifact removal on ChIP quality metrics in ChIP-seq and ChIP-exo data. 2014;5:75.

Chandler, J.W.J.P., cell environment. Auxin response factors. 2016;39(5):1014–1028.

Chaw, S.-M., et al. Dating the monocot–dicot divergence and the origin of core eudicots using whole chloroplast genomes. 2004;58(4):424–441.

Cheng, C., et al. Understanding transcriptional regulation by integrative analysis of transcription factor binding data. 2012;22(9):1658–1667.

Cochran, K., et al. Domain adaptive neural networks improve cross-species prediction of transcription factor binding. bioRxiv 2021.

Crisp, P.A., et al. Stable unmethylated DNA demarcates expressed genes and their cis-regulatory space in plant genomes. Proceedings of the National Academy of Sciences 2020;117(38):23991–24000.

Elkon, R. and Agami, R. Characterization of noncoding regulatory DNA in the human genome. Nature biotechnology 2017;35(8):732.

Epstein, D.J. Cis-regulatory mutations in human disease. Briefings in Functional Genomics and Proteomics 2009;8(4):310–316.

Estabrooks, A., Jo, T. and Japkowicz, N. A multiple resampling method for learning from imbalanced data sets. Computational intelligence 2004;20(1):18–36.

Freire-Rios, A., et al. Architecture of DNA elements mediating ARF transcription factor binding and auxin-responsive gene expression in Arabidopsis. 2020;117(39):24557–24566.

Galli, M., et al. The DNA binding landscape of the maize AUXIN RESPONSE FACTOR family. 2018;9(1):1–14.

Guilfoyle, T., et al. How does auxin turn on genes? 1998;118(2):341–347.

Guo, Y., Mahony, S. and Gifford, D.K. High resolution genome wide binding event finding and motif discovery reveals transcription factor spatial binding constraints. 2012.

Haberle, V. and Stark, A. Eukaryotic core promoters and the functional basis of transcription initiation. Nature reviews Molecular cell biology 2018;19(10):621–637.

Huang, D., et al. Identification of human silencers by correlating cross-tissue epigenetic profiles and gene expression. Genome research 2019;29(4):657–667.

Inukai, S., Kock, K.H. and Bulyk, M.L. Transcription factor–DNA binding: beyond binding site motifs. Current opinion in genetics & development 2017;43:110–119.

Kato, H., et al. Auxin-mediated transcriptional system with a minimal set of components is critical for morphogenesis through the life cycle in Marchantia polymorpha. 2015;11(5):e1005084.

Kato, H., et al. Design principles of a minimal auxin response system. 2020;6(5):473–482.

Kelley, D.R., Snoek, J. and Rinn, J.L.J.G.r. Basset: learning the regulatory code of the accessible genome with deep convolutional neural networks. 2016;26(7):990–999.

Langmead, B. and Salzberg, S.L.J.N.m. Fast gapped-read alignment with Bowtie 2. 2012;9(4):357–359.

Latchman, D.S. Transcription factors: an overview. The international journal of biochemistry & cell biology 1997;29(12):1305–1312.

Li, H., et al. The sequence alignment/map format and SAMtools. Bioinformatics 2009;25(16):2078–2079.

Li, S.-B., et al. A review of auxin response factors (ARFs) in plants. 2016;7:47.

Li, Y., Shi, W. and Wasserman, W.W.J.B.b. Genome-wide prediction of cis-regulatory regions using supervised deep learning methods. 2018;19(1):1–14.

Li, Y., Sun, G. and Zhu, Y. Data imbalance problem in text classification. In, 2010 Third International Symposium on Information Processing. IEEE; 2010. p. 301–305.

Lu, Z., et al. The prevalence, evolution and chromatin signatures of plant regulatory elements. Nature Plants 2019;5(12):1250–1259.

Martin, M. Cutadapt removes adapter sequences from high-throughput sequencing reads. EMBnet. journal 2011;17(1):10–12.

Mejía-Guerra, M.K. and Buckler, E.S.J.B.p.b. A k-mer grammar analysis to uncover maize regulatory architecture. 2019;19(1):1–17.

Mrázek, J. Finding sequence motifs in prokaryotic genomes—a brief practical guide for a microbiologist. Briefings in bioinformatics 2009;10(5):525–536.

Nakagawa, S., et al. DNA-binding specificity changes in the evolution of forkhead transcription factors. 2013;110(30):12349–12354.

O’Malley, R.C., et al. Cistrome and epicistrome features shape the regulatory DNA landscape. Cell 2016;165(5):1280–1292.

Oh, E., et al. Cell elongation is regulated through a central circuit of interacting transcription factors in the Arabidopsis hypocotyl. elife 2014;3:e03031.

Oka, R., et al. Genome-wide mapping of transcriptional enhancer candidates using DNA and chromatin features in maize. Genome biology 2017;18(1):1–24.

Ou, J., et al. motifStack for the analysis of transcription factor binding site evolution. 2018;15(1):8–9.

Park, P.J. ChIP–seq: advantages and challenges of a maturing technology. Nature reviews genetics 2009;10(10):669–680.

Park, P.J.J.N.r.g. ChIP–seq: advantages and challenges of a maturing technology. 2009;10(10):669–680.

Pennacchio, L.A., et al. Enhancers: five essential questions. Nature Reviews Genetics 2013;14(4):288–295.

Quang, D. and Xie, X. Dan Q: a hybrid convolutional and recurrent deep neural network for quantifying the function of DNA sequences. Nucleic acids research 2016;44(11):e107–e107.

Ramírez, F., et al. deepTools: a flexible platform for exploring deep-sequencing data. 2014;42(W1):W187–W191.

Ricci, W.A., et al. Widespread long-range cis-regulatory elements in the maize genome. Nature plants 2019;5(12):1237–1249.

Saeys, Y., Inza, I. and Larranaga, P. A review of feature selection techniques in bioinformatics. bioinformatics 2007;23(19):2507–2517.

Schmitz, R.J., et al. Quality control and evaluation of plant epigenomics data. The Plant Cell 2021.

Sielemann, J., et al. Local DNA shape is a general principle of transcription factor binding specificity in Arabidopsis thaliana. 2021;12(1):1–8.

Siggers, T. and Gordân, R.J.N.a.r. Protein–DNA binding: complexities and multi-protein codes. 2014;42(4):2099–2111.

Stormo, G.D., et al. Use of the ‘Perceptron’algorithm to distinguish translational initiation sites in E. coli. 1982;10(9):2997–3011.

Studer, A., et al. Identification of a functional transposon insertion in the maize domestication gene tb1. Nature genetics 2011;43(11):1160–1163.

Sun, Y., Wong, A.K. and Kamel, M.S. Classification of imbalanced data: A review. International journal of pattern recognition and artificial intelligence 2009;23(04):687–719.

Tiwari, S.B., Hagen, G. and Guilfoyle, T.J.T.P.C. The roles of auxin response factor domains in auxin-responsive transcription. 2003;15(2):533–543.

Ulmasov, T., Hagen, G. and Guilfoyle, T.J.J.P.o.t.N.A.o.S. Activation and repression of transcription by auxin-response factors. 1999;96(10):5844–5849.

Wang, L., et al. Altered chromatin architecture and gene expression during polyploidization and domestication of soybean. The Plant Cell 2021.

Weber, B., et al. Plant enhancers: a call for discovery. Trends in plant science 2016;21(11):974–987.

Wei, H., et al. Genome-Wide Identification of the ARF Gene Family and ARF3 Target Genes Regulating Ovary Initiation in Hazel via ChIP Sequencing. Frontiers in plant science 2021:1647.

Weirauch, M.T., et al. Determination and inference of eukaryotic transcription factor sequence specificity. 2014;158(6):1431–1443.

Xi, Y. and Li, W. BSMAP: whole genome bisulfite sequence MAPping program. BMC bioinformatics 2009;10(1):1–9.

Yang, A., et al. Review on the Application of Machine Learning Algorithms in the Sequence Data Mining of DNA. Frontiers in Bioengineering and Biotechnology 2020;8:1032.

Zamanighomi, M., et al. Predicting transcription factor binding motifs from DNA-binding domains, chromatin accessibility and gene expression data. Nucleic acids research 2017;45(10):5666–5677.

Zhang, Y., Jin, R. and Zhou, Z.-H. Understanding bag-of-words model: a statistical framework. International Journal of Machine Learning and Cybernetics 2010;1(1-4):43–52.

