## Supplemental Tables and Figures for "Identifying transcription factor-DNA interactions using machine learning"

| | Whole genome | | | UMR | | |
|---|---|---|---|---|---|---|
| ARF | # of peaks | bp of peaks | % in genome | # of peaks | bp of peaks | % in UMR |
| ARF4 | 32,115 | 6,423,000 | 0.30 | 13,563 | 2,726,163 | 2.21 |
| ARF16 | 45,107 | 9,021,400 | 0.43 | 16,639 | 3,344,439 | 2.72 |
| ARF18 | 8,938 | 1,787,600 | 0.08 | 4,102 | 824,502 | 0.67 |
| ARF27 | 63,865 | 12,773,000 | 0.61 | 30,171 | 6,064,371 | 4.92 |
| ARF29 | 15,718 | 3,143,600 | 0.15 | 7,171 | 1,441,371 | 1.17 |
| ARF34 | 117,709 | 23,541,800 | 1.12 | 31,570 | 6,345,570 | 5.15 |
| ARF7 | 32,099 | 6,419,800 | 0.30 | 4,031 | 810,231 | 0.66 |
| ARF10 | 35,540 | 7,108,000 | 0.34 | 9,154 | 1,839,954 | 1.49 |
| ARF14 | 29,142 | 5,828,400 | 0.28 | 3,621 | 727,821 | 0.59 |
| ARF25 | 40,468 | 8,093,600 | 0.38 | 8,584 | 1,725,384 | 1.40 |
| ARF36 | 22,414 | 4,482,800 | 0.21 | 2,979 | 598,779 | 0.49 |
| ARF39 | 10,966 | 2,193,200 | 0.10 | 2,211 | 444,411 | 0.36 |
| Average | 37,840.08 | 7,568,017 | 0.35 | 11,149.66 | 2,241,083 | 1.82 |

**Supplementary Table 1.** The number of peaks identified from DAP-seq data in maize and the proportion of the genome they occupy and the proportion that overlap with a UMR.

| Accuracy | | | | | | |
|---|---|---|---|---|---|---|
| | 75bp | 100bp | 125bp | 150bp | 175bp | Average |
| ARF non-bound | 80.37±0.6 | 86.3±0.66 | 88.1±1.07 | 89.18±1.06 | 90.15±0.73 | 80.37±0.6 |
| ARF-bound | 77.03±1.59 | 82.16±2.17 | 86.14±1.9 | 88.1±1.42 | 90.7±0.81 | 77.03±1.59 |
| **False Positive Rate (FPR)** | | | | | | |
| | 75bp | 100bp | 125bp | 150bp | 175bp | Average |
| ARF non-bound | 10.32±0.33 | 7.7±0.4 | 7.24±0.7 | 5.35±0.61 | 3.04±0.15 | 6.73 |
| ARF bound | 34.97±3.49 | 26.58±3.71 | 21.66±2.94 | 20.77±2.59 | 19.47±1.95 | 24.69 |
| **False Negative Rate (FNR)** | | | | | | |
| | 75bp | 100bp | 125bp | 150bp | 175bp | Average |
| ARF non-bound | 33.57±1.06 | 22.67±1.07 | 18.87±1.64 | 19.01±1.79 | 20.03±2.03 | 22.83 |
| ARF bound | 14.96±0.38 | 11.99±1.17 | 8.64±1.21 | 5.97±0.66 | 2.51±0.16 | 8.81 |

**Supplementary Table 2.** Accuracy, FPR and FNR based on the combinations of bin size and labelling method implemented. The model was validated by 5-fold cross-validation and repeated three times. Values represent the mean of accuracy ± variance.

| | | # of ARF-non-bound regions | # of ARF- bound regions | Ratio |
|---|---|---|---|---|
| Clade A ARF | ARF4 | 916,434 | 8,347 | 1:109.79 |
| | ARF16 | 914,743 | 10,038 | 1:91.12 |
| | ARF18 | 922,241 | 2,540 | 1:363.08 |
| | ARF27 | 906,168 | 18,614 | 1:48.68 |
| | ARF29 | 920,424 | 4,357 | 1:211.25 |
| | ARF34 | 905,355 | 19,426 | 1:46.60 |
| Clade B ARF | ARF7 | 922,314 | 2,467 | 1:373.86 |
| | ARF10 | 919,145 | 5,636 | 1:163.08 |
| | ARF14 | 922,600 | 2,181 | 1:423.01 |
| | ARF25 | 919,527 | 5,254 | 1:175.01 |
| | ARF36 | 922,956 | 1,825 | 1:505.72 |
| | ARF39 | 923,470 | 1,311 | 1:704.40 |
| | **Average** | **917,948.08** | **6,833** | **1:134** |

**Supplementary Table 3.** The number ARF-bound and ARF-non-bound regions used as classifications for the 12 ARFs evaluated in the maize genome. These results show the number of unbound DNA is significantly higher than that of bound DNA, which leads to imbalanced data issues.

| | Accuracy | FPR | FNR |
|---|---|---|---|
| 5-mer | 80.0±0.0 | 11.0±0.0 | 36.33±0.57 |
| 6-mer | 82.0±0.0 | 9.0±0.0 | 35.0±0.0 |
| 7-mer | 85.0±0.0 | 5.0±0.0 | 31.0±0.0 |
| 8-mer | 87.0±0.0 | 1.0±0.0 | 34.0±0.0 |
| 9-mer | 83.0±0.0 | 0.0±0.0 | 47.0±0.0 |

**Supplementary Table 4.** The performance of classifications based on different lengths of k-mers.

| ARF | Total # of reads | # of aligned reads (Ratio of aligned reads) |
|---|---|---|
| ARF4 | 13,511,580 | 13,021,975 (96.38%) |
| ARF16 | 8,478,227 | 8,131,473 (95.91%) |
| ARF18 | 4,026,230 | 3,879,889 (96.37%) |
| ARF27 | 12,791,259 | 12,259,171 (95.84%) |
| ARF29 | 12,840,351 | 12,350,446 (96.18%) |
| ARF34 | 11,102,829 | 10,706,386 (96.43%) |
| ARF7 | 9,293,703 | 8,907,608 (95.85%) |
| ARF10 | 6,427,418 | 6,170,068 (96.00%) |
| ARF14 | 11,153,468 | 10,710,799 (96.03%) |
| ARF25 | 10,611,447 | 10,099,664 (95.18%) |
| ARF36 | 9,391,176 | 9,033,961 (96.20%) |
| ARF39 | 10,147,655 | 9,681,922 (95.41%) |

**Supplementary Table 5.** Library size and alignment rate of DAP-seq for 12 maize ARFs screened against the soybean genome.

| ARF | Whole genome # of peaks | bp of peaks | % in genome | UMR # of peaks | bp of peaks | % in UMR |
|---|---|---|---|---|---|---|
| ZmARF4 | 19,478 | 3,895,600 | 0.41 | 7,651 | 1,537,851 | 0.51 |
| ZmARF16 | 16,964 | 3,392,800 | 0.35 | 6,282 | 1,262,682 | 0.42 |
| ZmARF18 | 1,106 | 221,200 | 0.02 | 455 | 91,455 | 0.03 |
| ZmARF27 | 10,248 | 2,049,600 | 0.21 | 4,471 | 898,671 | 0.30 |
| ZmARF29 | 5,395 | 1,079,000 | 0.11 | 2,513 | 505,113 | 0.17 |
| ZmARF34 | 35,715 | 7,143,000 | 0.74 | 12,407 | 2,493,807 | 0.82 |
| ZmARF7 | 20,802 | 4,160,400 | 0.43 | 6,112 | 1,228,512 | 0.41 |
| ZmARF10 | 12,286 | 2,457,200 | 0.26 | 4,393 | 882,993 | 0.29 |
| ZmARF14 | 9,542 | 1,908,400 | 0.20 | 2,940 | 590,940 | 0.20 |
| ZmARF25 | 19,058 | 3,811,600 | 0.40 | 6,957 | 1,398,357 | 0.46 |
| ZmARF36 | 22,516 | 4,503,200 | 0.47 | 6,938 | 1,394,538 | 0.46 |
| ZmARF39 | 4,210 | 842,000 | 0.09 | 1,541 | 309,741 | 0.10 |
| **Average** | **147,76.66** | **2,955,333** | **0.3** | **5,222** | **1,049,555** | **0.35** |

**Supplementary Table 6.** The number of DAP-seq peaks across the genome and within soybean UMRs.

| | | # of ARF-non-bound regions | # of ARF- bound regions | Ratio |
|---|---|---|---|---|
| Clade A ARF | ZmARF4 | 2,324,369 | 4,672 | 1:497.51 |
| | ZmARF16 | 2,325,206 | 3,835 | 1:606.31 |
| | ZmARF18 | 2,328,772 | 269 | 1:8,657.14 |
| | ZmARF27 | 2,326,307 | 2,734 | 1:850.88 |
| | ZmARF29 | 2,327,512 | 1,529 | 1:1,522.24 |
| | ZmARF34 | 2,321,402 | 7,639 | 1:303.88 |
| Clade B ARF | ZmARF7 | 2,325,264 | 3,777 | 1:615.63 |
| | ZmARF10 | 2,326,386 | 2,655 | 1:876.22 |
| | ZmARF14 | 2,327,267 | 1,774 | 1:1,311.87 |
| | ZmARF25 | 2,324,843 | 4,198 | 1:553.79 |
| | ZmARF36 | 2,324,806 | 4,235 | 1:548.95 |
| | ZmARF39 | 2,328,132 | 909 | 1:2,561.20 |
| | **Average** | **2,325,855.5** | **3,185.5** | **1:1,575.47** |

**Supplementary Table 7.** The number of events in each class used for the classification of the DAP-seq data from 12 maize ARFs screened against the soybean genome.

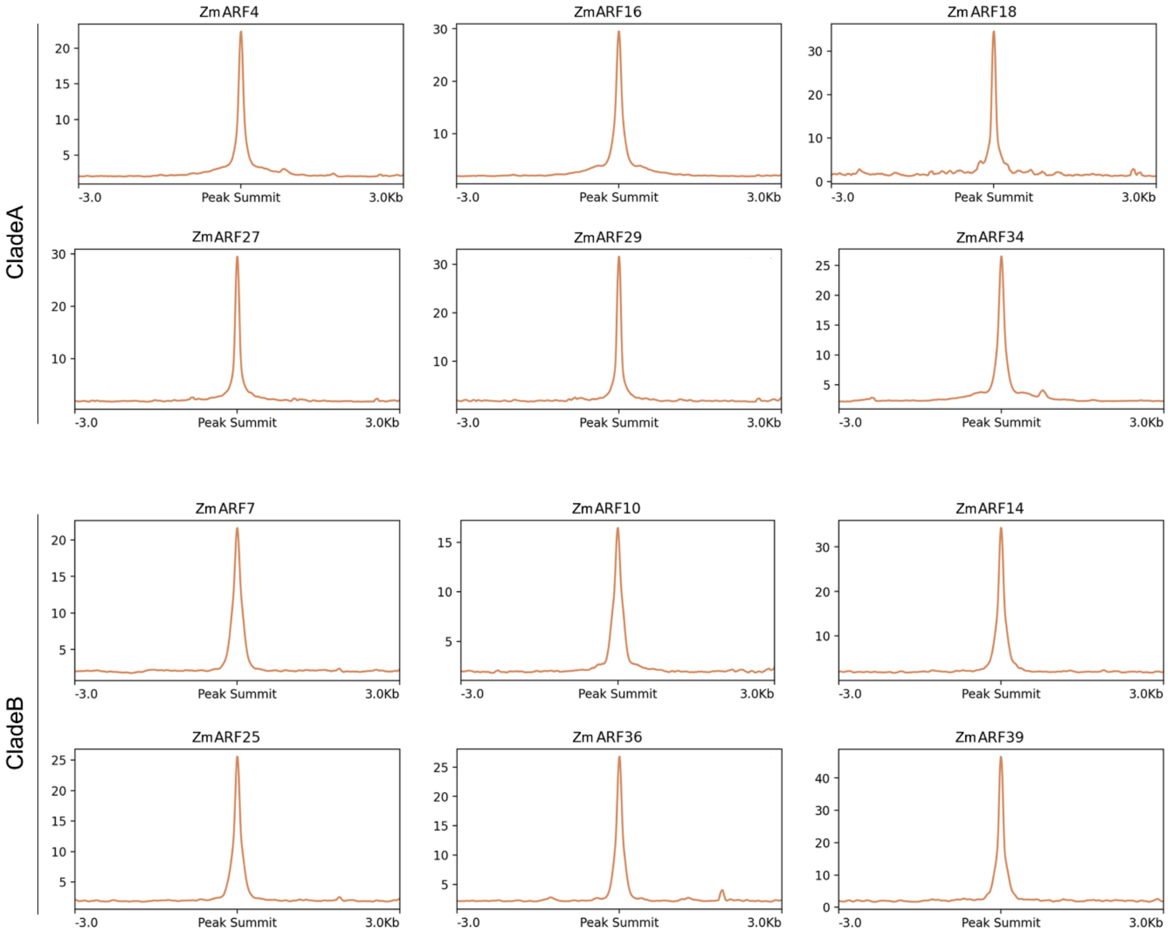


**Supplementary Figure 1. Distance to peak summit for DAP-seq.** Metaplots show the abundance around DAP-seq peaks for 12 members of ZmARFs in Soybean.

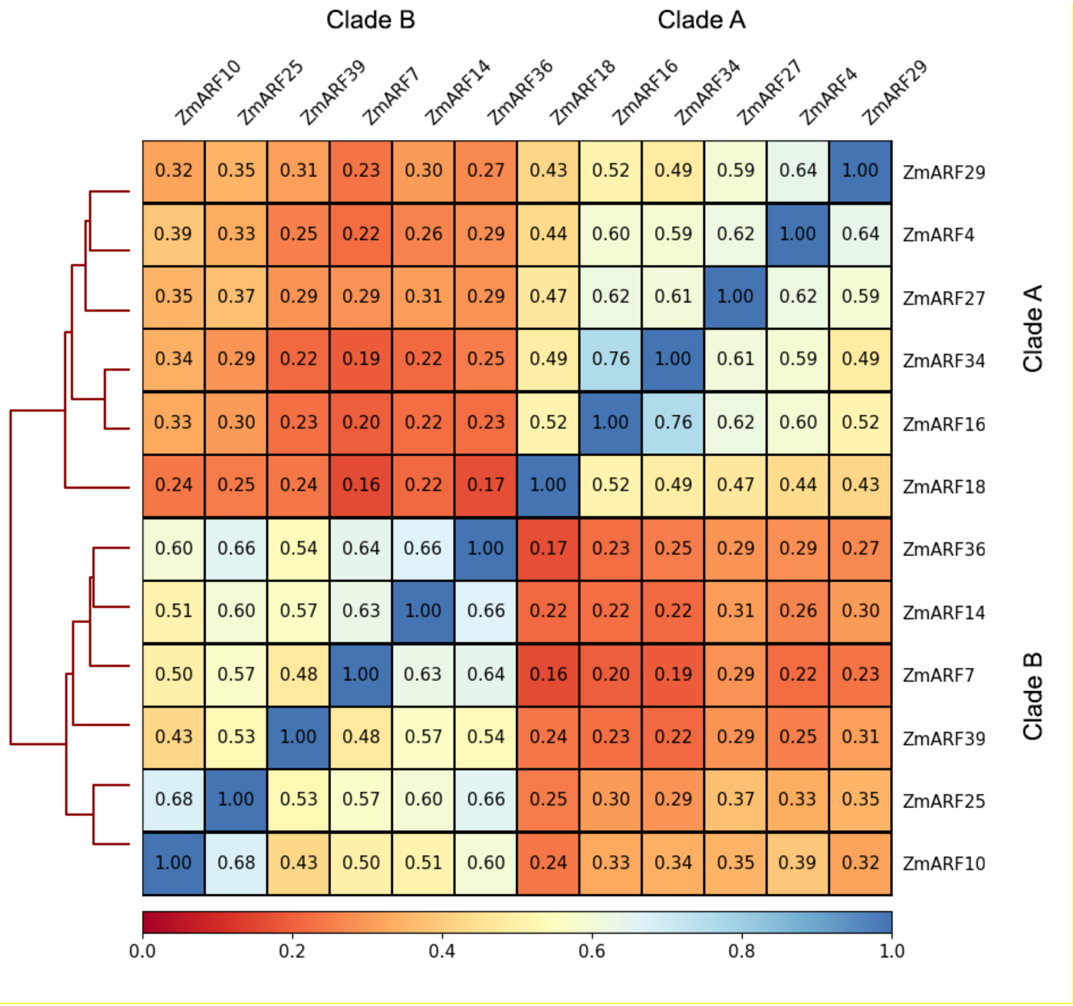


**Supplementary Figure 2.** Pairwise Pearson correlation analysis for soybean ZmARF binding events. All samples clustered according to their clade A or B phylogenetic classification.

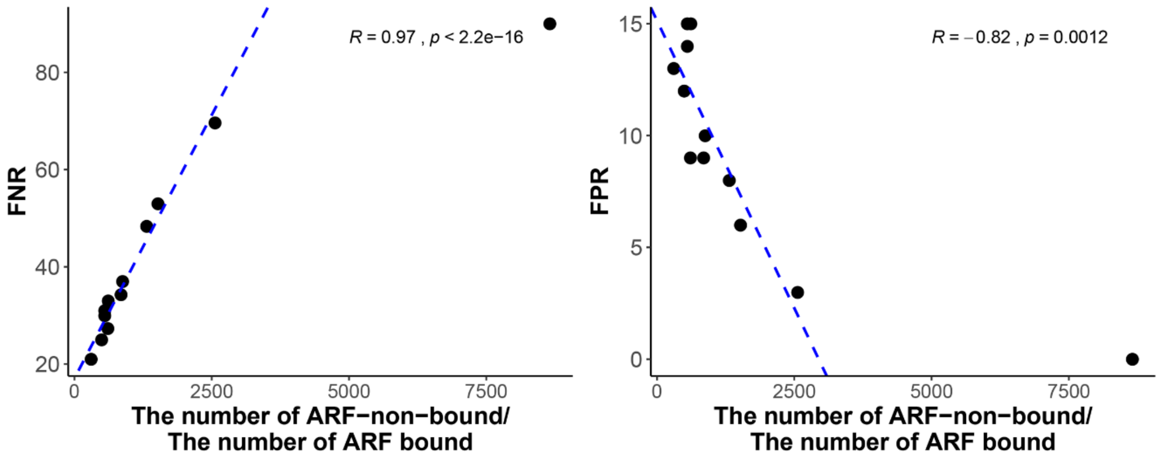


**Supplementary Figure 3.** The correlation between the ratio of imbalanced data from soybean ARF-bound regions with the FNR or the FPR. Each dots represents each of the 12 ZmARFs tested in this study. *R* is Spearman rank correlation coefficient.
